# A language model assistant for biocatalysis

**DOI:** 10.1101/2024.11.15.623739

**Authors:** Yves Gaetan Nana Teukam, Francesca Grisoni, Matteo Manica

## Abstract

Language model assistants have transformed how researchers interact with computational tools, offering unprecedented capabilities in understanding and generating complex scientific queries. We introduce a language model assistant for biocatalysis (LM-ABC), a computational tool designed to streamline workflows in enzyme engineering research. LM-ABC integrates a large language model with domain-specific modules to facilitate biocatalysis research through natural language inputs. Its architecture employs the Reasoning and Acting (ReACT) framework for dynamic tool selection and chaining, enabling functionalities like binding site extraction and molecular dynamics simulations. LM-ABC can interpret and process user queries in the form of natural language, and interface with existing computational resources to generate relevant results for enzyme engineering. Additionally, LM-ABC is available via both command-line and web-based interfaces, which lowers the barriers for its usage and integration in various disciplines. Provided as open-source software, the LM-ABC contributes to the application of language models in computational biology, potentially accelerating enzyme engineering research processes.

## Introduction

The rise of language model assistants (also called ‘agents’) marks a significant advancement across multiple scientific disciplines (32; 13; 6; 16; 11). By integrating large language models like GPT (7) and LLaMA (29) with task-specific tools, these assistants can interpret complex queries, reason about problems, and execute multi-step tasks in response to natural language inputs. Language model agents have fundamentally altered how researchers engage with computational tools (18; 10), enabling the automation of complex analytical processes, guiding experimental designs, optimizing workflows, and accelerating data-driven discoveries.

Language model assistants have found applications across various domains of biology and chemistry with unprecedented versatility. For instance, ChemCrow (19), IBM’s ChatChem (14), and ChatMOF (15), streamline chemistry and material sciences by integrating tools for cheminformatics, reaction optimization and molecular property predictions. In proteomics, assistants like ProtAgent (12) are employed for protein information retrieval and structure generation.

**Figure 1:**
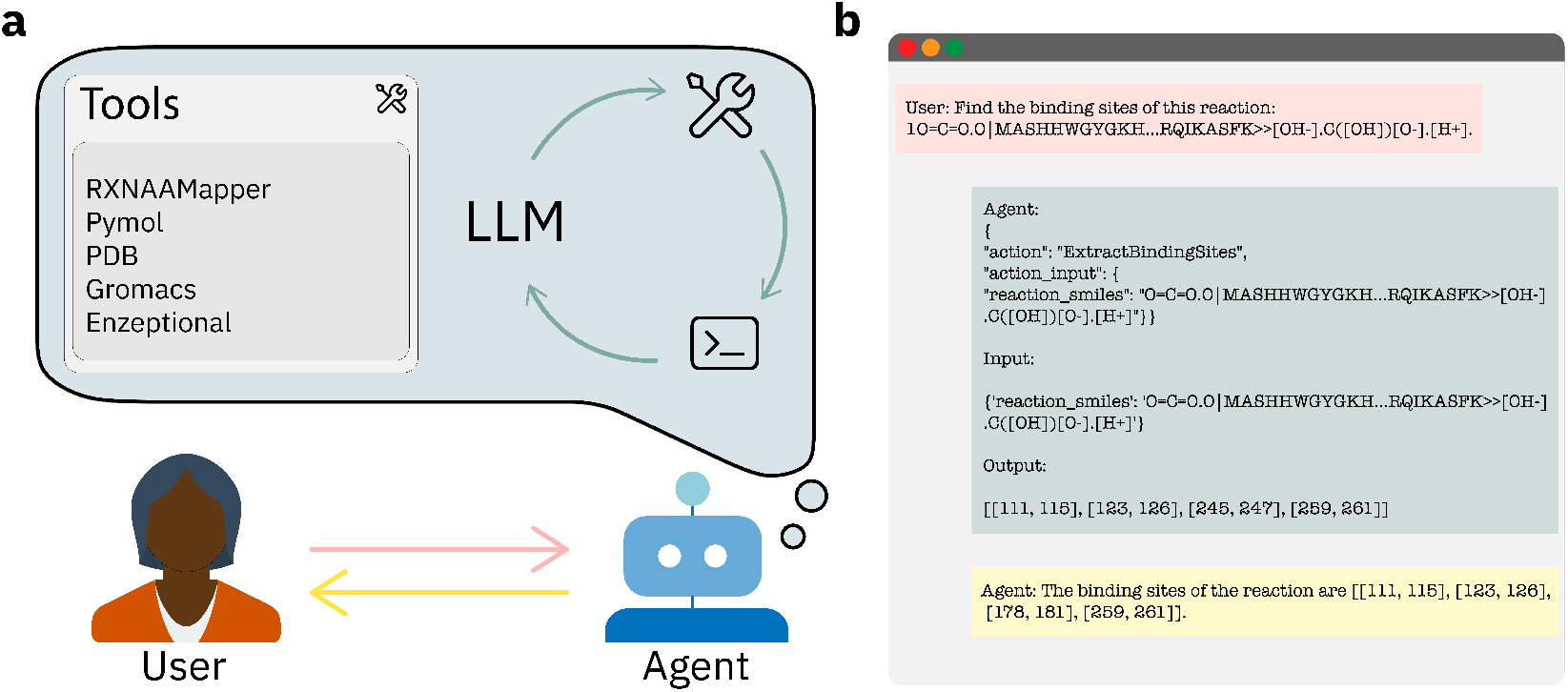
Language Model Assistant for BioCatalysis (LM-ABC). (a) The agent, through a large language model (LLM) core processes natural language inputs from the user, and it selects specialized tools to address the required tasks. (b) Example of interaction between the user and the agent, revealing the agent’s internal reasoning. Here, the agent employs the `ExtractBindingSites` tool to analyse a reaction SMILES string.

Biocatalysis is a time- and resource-intensive endeavour (4; 22), which could benefit enormously from language models agents. In fact, biocatalysis demands the integration of precise, context-specific computational tools to design experiments, optimize enzymes, and navigate the complex landscape of enzyme function and structure. Despite this potential, the field of biocatalysis lags behind other domains of chemistry when it comes to tailored assistants specifically designed for enzyme engineering. To date, computational methods for biocatalysis (*e.g*. molecular dynamics simulations, structure prediction, and sequence alignment), require manual integration, expert interpretation, and extensive computational resources (28). Such a ‘fragmented’ approach slows down progress in biocatalysis, underscoring the need for holistic computational solutions that are broadly accessible.

Here, we introduce the language model assistant for biocatalysis (LM-ABC), which fills this gap by introducing a language model assistant designed to support biocatalysis and bioinformatics tasks. Unlike existing tools that operate in isolation or require specialized knowledge, this assistant integrates a comprehensive set of functionalities into a unified, interactive platform. Key features include the ability to extract binding sites from enzyme sequences, optimize enzyme variants for improved catalytic activity, perform in silico mutagenesis, and conduct molecular dynamics simulations—all accessible through an intuitive, language-guided interface. This integration not only streamlines computational workflows but also empowers researchers to perform complex analyses without deep computational expertise.

Thanks to its specialized framework integrating bioinformatics tools, LM-ABC constitutes a significant advancement for biocatalysis research. By enabling a holistic approach to enzyme design, enhanced by the capabilities of large language models, we expect LM-ABC to accelerate the pace of enzyme optimization and molecular design, ultimately contributing to pushing the boundaries of what is possible in biocatalysis research.

## Biocatalysis Assistant

### System Overview

LM-ABC is built on a modular architecture that integrates advanced natural language processing techniques, dynamic tool management, and an automated execution pipeline. At its core, the assistant employs a large language model (LLM) that can be integrated with various providers. The system supports a wide range of modern LLMs, including OpenAI’s GPT-3 and GPT-4 series (7; 2), Anthropic’s Claude models, Hugging Face’s opensource models, and local deployments through Ollama (21). The system uses the Llama 70B instruct model as its default model, but users can explicitly specify their preferred model and provider combination when needed. This LLM functions as the primary interface, interpreting user inputs and translating them into executable workflows within the biocatalysis domain.

We implemented the assistant using the ReACT (Reasoning and Acting) framework, leveraging the langchain library (8). This framework combines the language understanding capabilities of transformer-based models with a decision-making module, enabling the assistant to dynamically adjust its reasoning process and action selection based on intermediate results and evolving task requirements.

The assistant interacts with various integrated tools, each accompanied by a comprehensive metadata describing its capabilities, input/output specifications, and appropriate usage contexts. This metadata is incorporated into the LLM’s system prompt, ensuring the model maintains an up-to-date understanding of available tools and their functions. This approach enhances the assistant’s ability to select and execute tools dynamically during complex workflows.

The LM-ABC incorporates several specialized tools designed to address key aspects of biocatalysis and bioinformatics analysis, detailed in the subsequent section. Once a tool is selected, the assistant oversees the execution process and output generation based on user inputs.

Results from the various tools are presented to the user in a clear and interpretable format including raw data, contextual insights, recommendations, and potential next steps derived from the language model’s interpretation. This final layer, combined with the overall system design, facilitates a seamless and intuitive user experience.

### Integrated Tools

The LM-ABC incorporates a diverse set of specialized tools, each addressing specific challenges in biocatalysis and bioinformatics. These tools form a cohesive ecosystem, automating critical steps in enzyme analysis and design to enhance the efficiency and accuracy of biocatalytic research.

#### Reaction parsing

Analysis of biochemical reactions requires deconstructing complex reaction strings into their constituent elements, including reactants, amino acid sequences, and products. This parsing is achieved through the `GetElementsOfReaction` tool, which processes reactions expressed in SMILES (Simplified Molecular Input Line Entry Systems) format (30).

#### Binding site extraction

Understanding enzyme functionality requires identifying key sites that can be targeted for mutations to enhance catalytic activity or optimize user-specified fitness functions. This critical analysis is performed by the `ExtractBindingSites` tool, which leverages RXNAAMapper (26) to extract binding sites from reaction SMILES strings.

#### Optimizing sequences

The optimization of enzyme sequences for biocatalytic reactions is crucial for improving their performance. This optimization process is handled by the `OptimizeEnzymeSequences` tool, built on the Enzeptional framework (27). It enables multiple optimization iterations based on substrate and product SMILES, featuring customizable scoring models and interval-specific mutations, ultimately producing a ranked list of optimized sequences for experimental validation.

#### Retrieving similar sequences

Protein homology and function analysis often requires identifying sequences similar to a given query. This search functionality is implemented through the `Blastp` tool, which uses NCBI’s Basic Local Alignment Search Tool for Proteins (3; 31). The tool provides comprehensive output including aligned sequences, descriptions, and statistical data, with customizable search parameters.

#### Retrieving protein structures

Access to threedimensional protein structures is essential for structural analysis and modeling. This functionality is provided by two complementary tools: `FindPDBStructure` and `DownloadPDBStructure`, which interface with the Protein Data Bank (PDB) (5) through the RCSB Search API (23) for structure identification and retrieval.

#### Targeted mutations

Protein structure modification to match specified target sequences is a crucial step in protein engineering. This transformation is accomplished through the `Mutagenesis` tool, which utilizes PyMOL (24) for mutation implementation and can optionally perform structural impact analyses such as RMSD calculations.

#### Molecular dynamics

Comprehensive simulation of protein dynamics requires multiple stages of equilibration and analysis. The `MDSimulation` tool automates this process using GROMACS (1), handling the setup and execution of standard MD simulation stages including Minimization, NVT (constant Number, Volume, Temperature) equilibration, and NPT (constant Number, Pressure, Temperature) equilibration.

### Interface

The LM-ABC offers two user interfaces: a Command Line Interface (CLI) and a web-based Streamlit application. These interfaces cater to different user preferences and use cases, providing flexibility in how researchers interact with the assistant.

#### Command Line Interface (CLI)

The CLI provides a traditional text-based interface suitable for terminal-based operations or integration into existing commandline workflows. The CLI can be accessed executing the command `lmabc` in the terminal following installation via Poetry (Eustace and The Poetry contributors).

#### Web Application

A web-based application, built on Streamlit, provides a graphical interface through a browser. The application be launched by executing the command `lmabc-app` in the terminal environment.

Both interfaces interact with the same underlying `BiocatalysisAssistant` class, ensuring consistency in functionality and results. This dual-interface approach caters to a wide range of user preferences and technical backgrounds, making the LM-ABC accessible to both command-line enthusiasts and those who prefer graphical interfaces.

### Use Case: Optimization of an Enzyme for Enhanced Catalytic Activity

To demonstrate the capabilities of our assistant, we present a use case focused on the optimization of the human Carbonic Anhydrase II (CA II). Starting from a substrate (CO expressed as O=C=O.O in SMILES notation) and the CA II protein sequence (retrieved from UniProt using the accession ID P00918), and the expected products (bicarbonate and proton, expressed as [OH-].C([OH])[O].[H+] in SMILES notation), we show how the assistant integrates multiple tools to streamline a complex workflow in enzyme engineering.

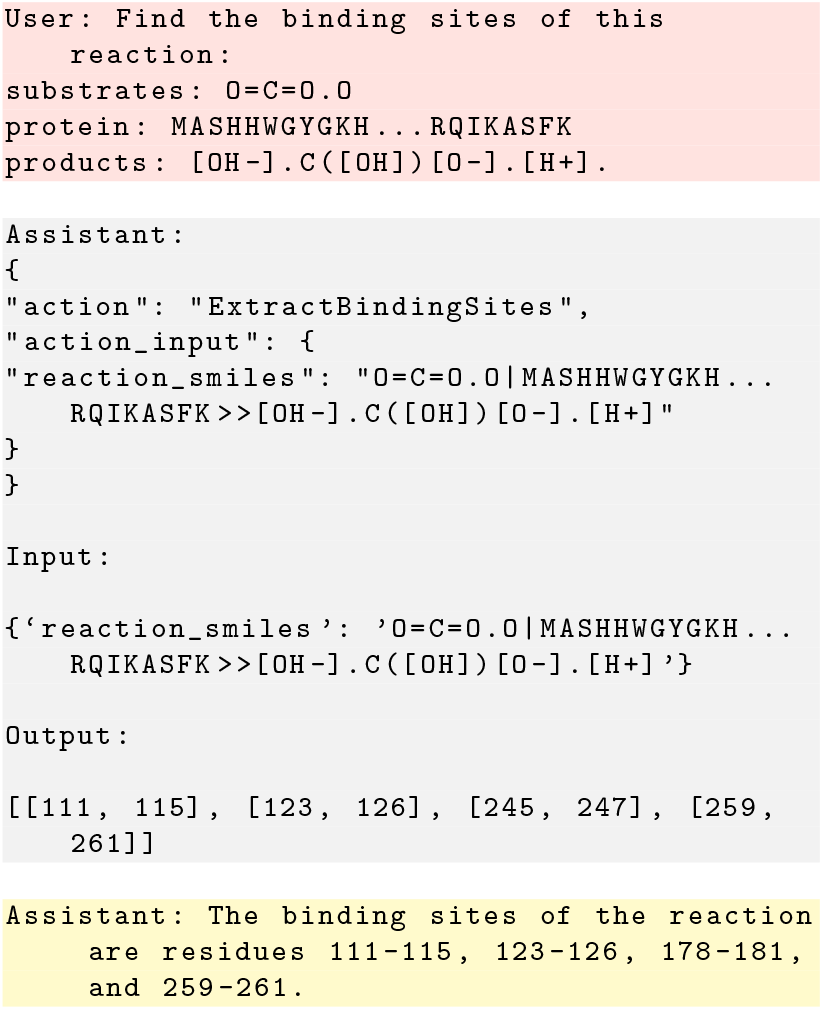

This interaction shows the assistant’s reasoning process and tool selection capability. The assistant uses a `CustomJSONAssistantOutputParser`, derived from Langchain’s `JSONAssistantOutputParser`, to process the language model’s outputs. This parser handles both structured `JSON` responses and raw text, ensuring robust interpretation of the model’s output.

The assistant’s response is formatted in `JSON`, adhering to the structure defined in the system prompt:

~~~
{
“ action “: “ ExtractBindingSites “,
“ action_input “: {
“ reaction_smiles “: “O=C=O.O| MASHHWGYGK
    H… RQIKASFK >>[OH−].C([OH])[O−].[H+]“
}
}
~~~

This format clearly distinguishes between the chosen action (tool selection) and the input parameters. The `ExtractBindingSites` tool, utilizing `RXNAAMapper`, processes the reaction SMILES string to identify potential binding sites within the enzyme sequence.

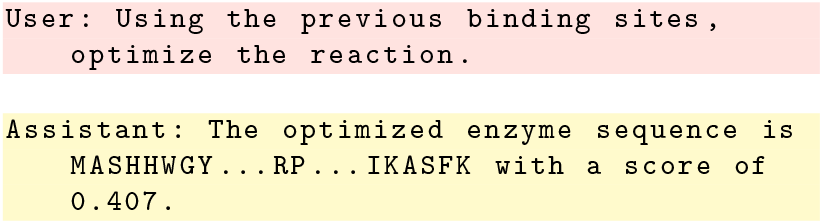

This step highlights the assistant’s memory capabilities. Using `ConversationBufferMemory`, the assistant maintains context across multiple interactions, crucial for handling complex, multi-step tasks in enzyme optimization. This memory system ensures coherence in extended biocatalysis workflows.

### Additional Resources

LM-ABC is provided as an open-source software on GitHub. Installation instructions, system requirements, and additional documentation are available on GitHub at the following URL: https://github.com/GT4SD/lm-assistant-for-biocatalysis

## Acknowledgements

Y.G.N.T would like to thank Oliver Schilter for their helpful suggestions when preparing this library.

## Supplementary Material

### Use Case: Carbonic Anhydrase II Optimization

Human Carbonic Anhydrase II (CA II; EC 4.2.1.1) is a well-studied zinc metalloenzyme that catalyzes the reversible hydration of carbon dioxide to bicarbonate and protons (17). This enzyme plays crucial roles in various physiological processes, including pH regulation, CO_2_ transport, and ion exchange (25). Given its importance in both biological systems and potential industrial applications, CA II serves as an ideal candidate for demonstrating the capabilities of our LM-ABC in enzyme optimization.

In this use case, we aim to optimize the catalytic efficiency of CA II for CO_2_ hydration using our integrated computational approach. The workflow encompasses several key steps: (1) identification of binding sites, (2) optimization of the enzyme sequence, (3) retrieval of the wild-type structure, (4) in silico mutagenesis, and (5) molecular dynamics simulation to validate the proposed optimizations.

#### 1. Binding Site Identification

We initiated the optimization process by identifying the binding sites of CA II using our `ExtractBindingSites` tool. The reaction SMILES string for CO_2_ hydration was input into the LM-ABC:

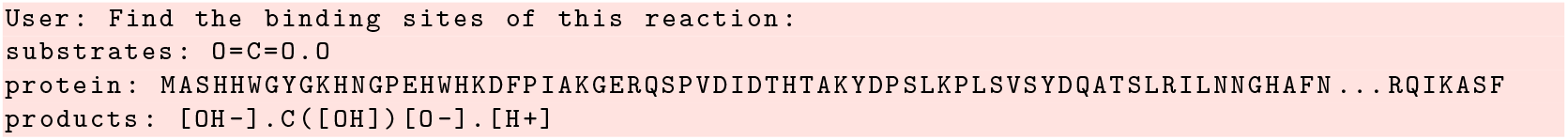

The LM-ABC provided the following output:

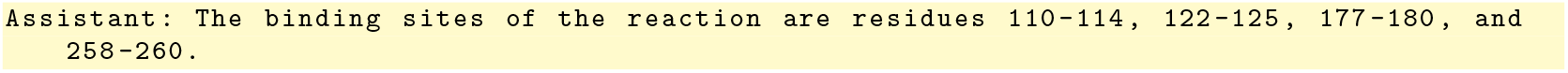

#### 2. Enzyme Sequence Optimization

Using the identified binding sites, we proceeded to optimize the CA II sequence for enhanced catalytic activity. The `OptimizeEnzymeSequences` tool was employed with the following input:

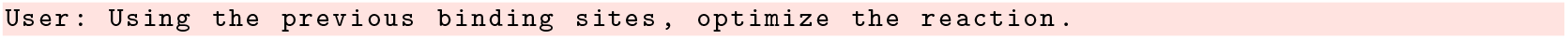

The LM-ABC generated a series of optimized sequences. The top 5 variants and their corresponding scores are:

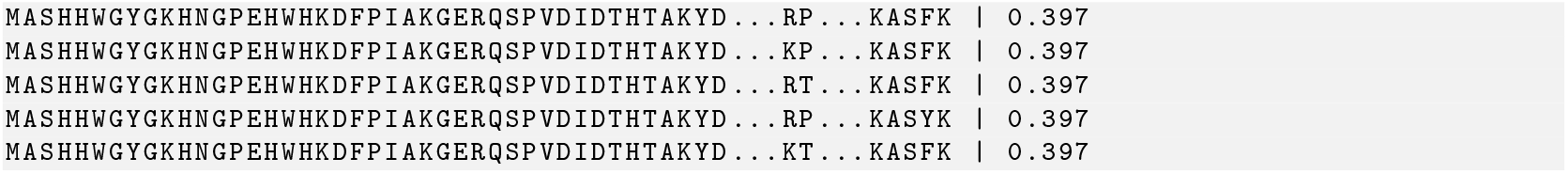

The top-scoring sequence achieved a score of 0.397.

#### 3. Retrieval of Wild-type Structure

To provide a structural context for the proposed mutations, we retrieved the wild-type CA II structure from the Protein Data Bank using the `FindPDBStructure` and `DownloadPDBStructure` tools:

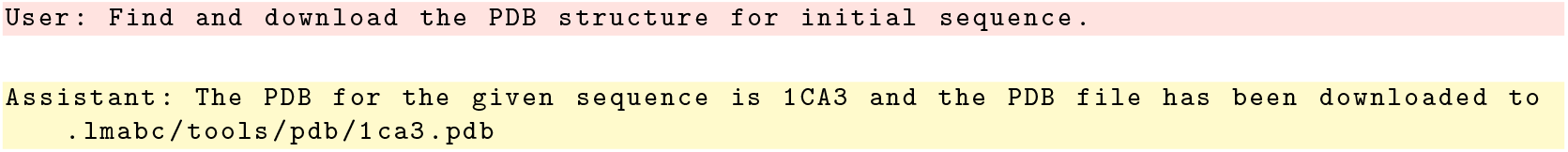

The assistant identified and retrieved PDB ID: 1CA3 (2.30 Å) crystal structure of human CA II (20).

#### 4. In Silico Mutagenesis

We performed in silico mutagenesis on the wild-type structure (1CA3) to generate a model of the top-ranked optimized sequence using the `Mutagenesis` tool:

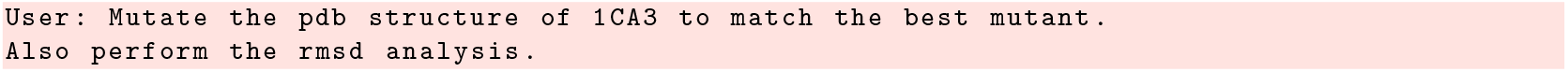

The LM-ABC executed the mutagenesis and provided a modified PDB file along with an RMSD calculation between the wild-type and mutant structures:

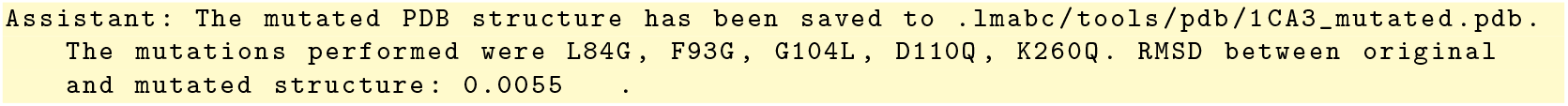

#### 5. Molecular Dynamics Simulation

To assess the dynamic behavior and stability of the optimized CA II variant, we conducted a molecular dynamics simulation using the `MDSimulation` tool:

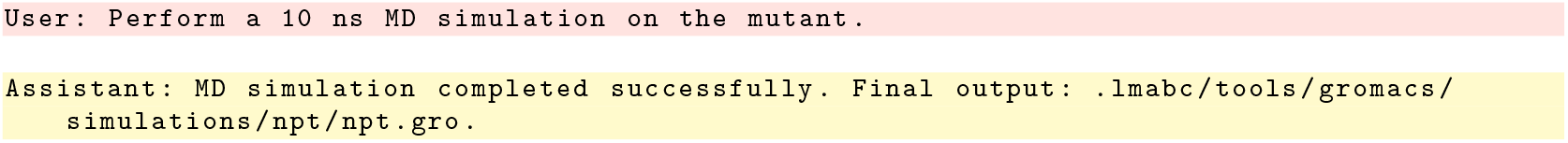

#### Web Interface Visualization

Figure S1 shows the web-based interface of LM-ABC, implemented using Streamlit. The interface features a clean, intuitive design with a sidebar containing quick access to tools, settings, documentation, and examples. The main panel displays the interaction area where users can input their queries in natural language. In this example, the interface shows a typical optimization workflow where the user requests sequence optimization for a CO_2_ hydration reaction. The assistant’s reasoning process and JSON-formatted tool selection are visible in the response, demonstrating the transparency of the decision-making process. The interface also includes proper citation information and GitHub repository access, making it a comprehensive tool for both academic and industrial users.

**Figure S1:**
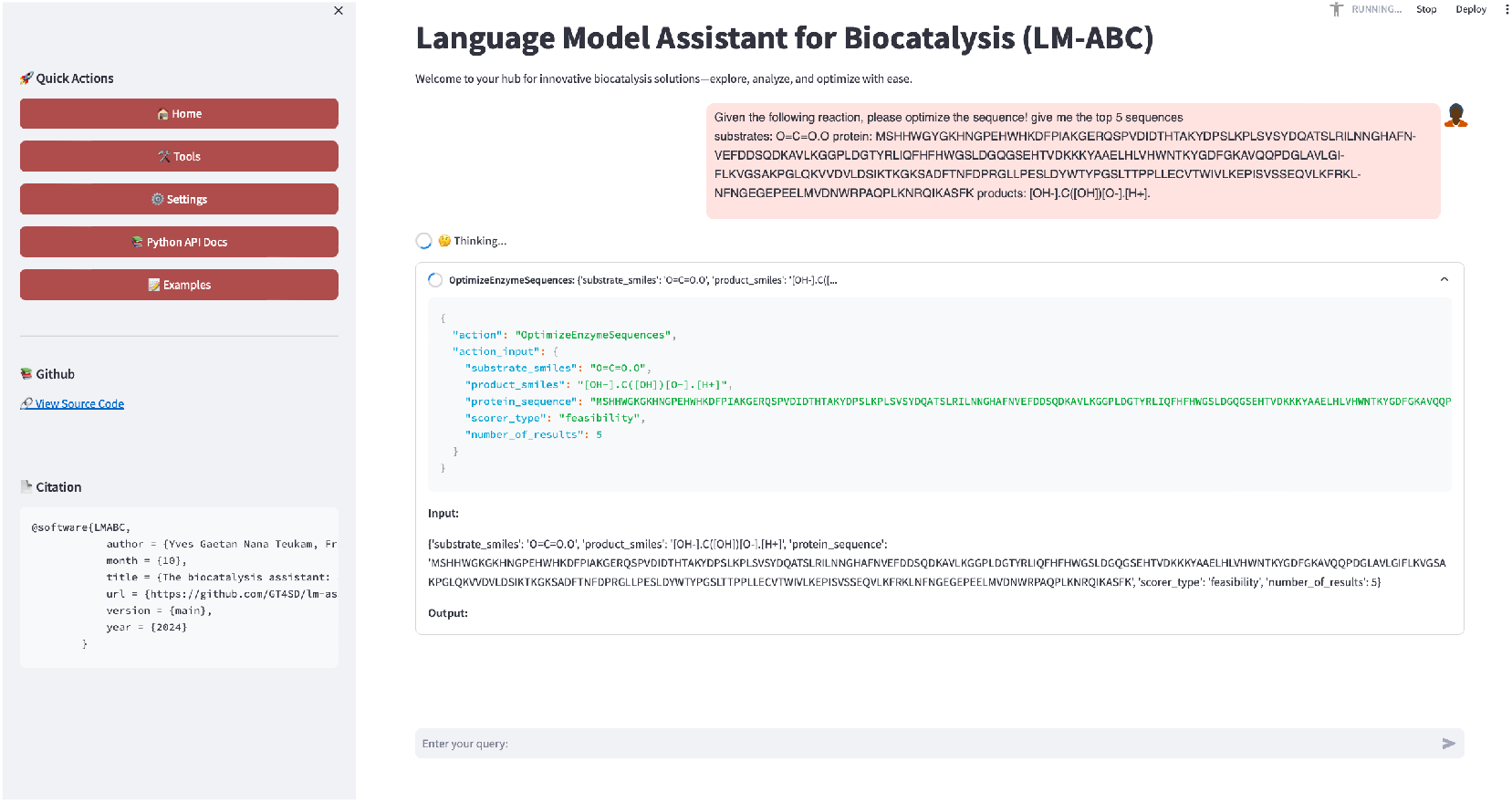
Web interface of LM-ABC. The interface shows the main components: navigation sidebar (left), query input area (bottom), and interaction display (center) with the assistant’s reasoning process visible.

## Discussion and Conclusions

This use case demonstrates the capability of the LM-ABC to perform a comprehensive computational workflow for enzyme optimization. The assistant successfully identified key catalytic residues in CA II, proposed sequence optimizations, and facilitated structural analysis of the mutations.

However, it is important to note that computational predictions require experimental validation. Future work should include the expression and characterization of the proposed CA II variants to confirm enhanced catalytic activity. Additionally, more extensive MD simulations and advanced sampling techniques could provide deeper insights into the structural and dynamic effects of the proposed mutations.

